# Emergent Stable Tissue Shapes from the Regulatory Feedback between Morphogens and Cell Growth

**DOI:** 10.1101/2025.02.16.638504

**Authors:** Bivash Kaity, Daniel Lobo

## Abstract

The development and regeneration of multicellular organisms require dynamic coordination between cellular behaviors and mechanochemical signals to achieve precise and stable tissue shapes. Plastic organisms, such as planarians, can regenerate, grow, and degrow as adults while maintaining precise whole-body and organ tissue shapes. However, the mechanisms underlying the pathways that coordinate and integrate these signals into the correct balance between cellular growth, mitosis, and apoptosis to form emergent target tissue shapes remain poorly understood. Here, we present a systematic theoretical study of the biological drivers controlling the feedback mechanisms between tissue growth and morphogen signaling. The approach is based on lattice-free, center-based simulations of cell size dynamics, mitosis, and apoptosis governed by both intercellular diffusible morphogen concentrations and mechanical stress between cells to drive their spatial organization. The results demonstrate how different morphogen properties and tissue mechanics form a feedback loop that is essential for the robust regulation of target tissue shapes. Furthermore, we show that stable tissue shapes can emerge from self-regulated patterning processes, such as Turing systems, controlling cellular growth dynamics. A stable feedback loop can form between the emergent morphogen patterns and the dynamics of cellular growth they regulate, as the tissue dynamics define the domain in which morphogens diffuse and hence pattern. Overall, this study highlights the essential role of the feedback loop between morphogen patterning and cellular growth in the regulation of tissue dynamics for stable shape formation. Moreover, this work establishes a framework for further experiments to understand the regulatory dynamics of whole-body development and regeneration using models with high spatiotemporal resolution.

**Significance:** Tight coordination and interpretation of the multitude of signals at different biological scales– from intracellular signals to mechanical interactions–are essential during the development and regeneration of multicellular organisms. In this work, we investigate the leading role of the feedback between mechanochemical signaling networks and tissue shape through cellular behaviors such as growth, proliferation, and apoptosis. This study demonstrates the interdependence between tissue growth and pattern formation mechanisms in the regulation of stable tissue shapes. Overall, this research provides novel mechanistic insights into the formation of tissue shapes through the regulatory feedback interaction between cell growth and patterning dynamics.

## 1. Introduction

Tissue growth, patterning, and differentiation are intricately interconnected processes that drive the development and regeneration of multicellular organisms. Crucially, these processes act at different biological scales to modulate fundamental cellular mechanisms (Bailles et al., 2022; Farahani and Nelson, 2022), including cell growth, proliferation, apoptosis, and migration (Collinet and Lecuit, 2021). In addition, biochemical signaling critically contributes to the emergence of shapes and forms by establishing the spatial distribution of morphogens, from organs (Eldred et al., 2017; Kudla et al., 2022; Ramezani et al., 2023; Salbreux et al., 2012), to organoids (Xue et al., 2025), to whole-body shapes (Lobo, 2022; Lobo et al., 2013; Lobo and Levin, 2015; Maroudas-Sacks et al., 2025; Roy et al., 2020). Indeed, plastic organisms such as planarians can precisely control their dynamic shape during development, regeneration, growth, and degrowth through the balanced regulation of mitosis, apoptosis, and differentiation by morphogens that react and diffuse within their dynamic tissue shapes (Adell et al., 2025; Davies et al., 2017; Emmons-Bell et al., 2015; Herath and Lobo, 2020; Ko et al., 2024; Lobo et al., 2012; Stückemann et al., 2017). Although it is known that the coordination and dynamic synchronization of cellular behaviors by morphogens regulate the size, shape, and function of an organism (Aegerter-Wilmsen et al., 2012; George et al., 2025; Heisenberg and Bellaïche, 2013), a mechanistic understanding of the feedback regulation between signaling and tissue growth to form stable target shapes during development and regeneration remains unclear.

The complex feedback loops between mechanochemical signaling and cellular mechanisms regulate patterning and growth, and hence, the emergence of shapes and forms. Dysregulation of this network may lead to abnormalities in tissue shape and function (Boulet et al., 2004; Lu et al., 2006; Thompson et al., 2016). Several mechanisms have been studied to explain the emergence of tissue shapes from mechanical interactions, including attractive forces due to cell-cell adhesion by extracellular matrix components (Ko and Lobo, 2019; Steinberg, 1958; Townes and Holtfreter, 1955), as well as repulsive forces due to their intrinsic elastic properties, which inhibit the deformation of cell morphology and can guide tissue patterning (Hagolani et al., 2019). The range of these interactions between cells plays a crucial role in their spatial organization and the resulting tissue shape (Vasilopoulos and Painter, 2016). For example, long-distance cell-cell interactions are essential for spreading epithelial cells (Salm and Pismen, 2012). In addition to mechanical interactions, chemical signals acting as morphogens significantly contribute to the regulation of cell growth, proliferation, and death, which are necessary for proper tissue morphogenesis (Aegerter-Wilmsen et al., 2012; Cano-Fernández et al., 2025; Economou et al., 2012; Kicheva and Briscoe, 2023). Morphogens can form self-regulated reaction-diffusion mechanisms to establish patterns of cell growth and differentiation towards stable tissue shapes (Green and Sharpe, 2015). Indeed, morphogen patterns based on Turing systems have been shown to drive digit formation (Raspopovic et al., 2014), feather patterning (Jung et al., 1998), ruggae in the mammalian palate (Economou et al., 2012), and scutes in turtle shells (Moustakas-Verho et al., 2014). However, understanding how these morphogen patterns drive the formation of specific and stable tissue shapes by precisely balancing cellular growth, mitosis, apoptosis, and differentiation remains challenging (Mateus et al., 2021).

To understand the feedback between cell behavior, signaling, and emergent tissue growth, mathematical and computational approaches have been proposed to model cell-level phenomena (Cockerell et al., 2023; Osborne et al., 2017; Runser et al., 2024; Sharpe, 2017). While continuous models have traditionally been employed to study tissue pattern formation, approaches based on off-lattice and cell-center models are especially suitable for simulating discrete cells that can grow, divide, and die to form free tissue growth dynamics and shapes without limited spatial resolution (Glen et al., 2019; Pleyer and Fleck, 2023; Van Liedekerke et al., 2015; West et al., 2023). In this manner, the cell positions are continuous in space and can dynamically change their neighbors over time. This exchange of neighbors contributes significantly to emergent behaviors, such as cell sorting, collective migration, and mechanical feedback (Mathias et al., 2020). Crucially, off-lattice models can also couple cellular and chemical sub-models by simulating morphogen reaction-diffusion systems in superimposed meshes derived from cellular positions (Kaul et al., 2023; Okuda et al., 2018a; Ramezani et al., 2023). This approach has been used to study the dynamics of cell pattern formation, including intestinal crypt patterning (Dunn et al., 2013; Van Leeuwen et al., 2009), early embryogenesis (Marin-Riera et al., 2015), zebrafish development (Delile et al., 2017), tumor growth (Ghaffarizadeh et al., 2018), mesenchymal and epithelial polarized cells (Germann et al., 2019), and Drosophila wing development (Muñoz-Nava et al., 2020). However, understanding the biological parameters that regulate the stability of the dynamic feedback between morphogen patterns and the tissue shape that they control remains an open problem (Mathias et al., 2022). It is essential to investigate which specific factors influencing and balancing cell growth, mitosis, and apoptosis can be modulated by organizers or self-regulating morphogenetic systems to produce distinct and stable tissue shapes.

Here, we present a systematic study of the ability of the feedback loop between morphogen patterning and tissue growth to form emergent stable tissue shapes using a mathematical and computational model of coupled cellular mechanics, signaling, and growth. The model is based on a lattice-free, agent-based framework that combines the roles of biochemical and mechanical signals at the cellular level to study emergent properties at the tissue level. The physical and adhesive properties of cells dictate their coordinated movement and arrangement. Morphogens that react at the cellular level diffuse at the tissue level to define specific spatial expression patterns, which in turn control cellular growth, mitosis, and apoptosis. As cells dynamically grow, divide, and die in response to morphogens, they define both the intercellular space for gene regulatory mechanisms and the tissue space in which intracellular signals diffuse, thereby closing the loop between tissue growth and morphogen patterning. Employing cell-center positions to calculate both mechanical and chemical interactions reduces the complexity and computational requirements of the model. We hypothesized that stable tissue shapes are influenced by two factors that directly affect cellular growth: the level of growth morphogens and the amount of mechanical stress caused by cell density. The results show that different biological parameters modulating the feedback between morphogens and tissue growth play different roles in regulating the stable shapes and sizes of emergent tissues. In addition, we demonstrate the ability of Turing reaction-diffusion systems to control the emergence of different stable tissue shapes because of the feedback mechanism between spot or stripe morphogen patterns that dynamically form as the tissue grows. Such feedback can reach equilibrium, resulting in a stable tissue shape. Overall, the proposed framework provides an understanding of the feedback between morphogen patterning and tissue-level growth through a systematic study of the biological parameters that regulate cell-level behaviors, such as motility, growth, division, degrowth, and death.

## 2. Methods

### 2.1. Cells

We define a model of an independent tissue, such as an organ or a whole body, made of a dynamic collection of cells, each considered a circular disc capable of migration, growth, division, and apoptosis. The disc center *x*_i_ represents the position of cell *i*, and the disk radius *r*_i_ defines its size. Therefore, a system of *N* cells at a given time is defined as a set of points {*x*_1_, *x*_2_,…, *x*_N_} and associated radii {*r*_1_, *r*_2_,…, *r*_N_}, with *x*_i_ ∈ ℝ^2^ and *r*_i_ ∈ ℝ. The movement of each cell *i* in the system is governed by Newton’s law of motion, such as

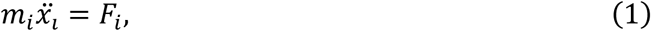

where *m*_i_ is the cell mass and *F*_i_ represents the sum of all forces acting on the cell. Each cell *i* is subjected to a viscous force *F*^vis^_i_ and an interaction force *F*^int^_i_, such that

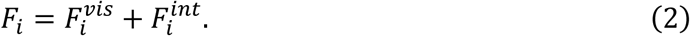

The viscous force occurs in the opposite direction of motion and is due to the interaction of a cell with the extracellular matrix and media, such that

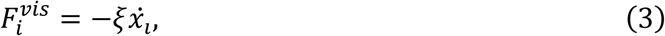

where ॐ is the viscosity coefficient. Since the acceleration due to the inertial force is negligible compared to the effect of the viscous force in the cell microenvironment (Purcell, 1977; Shu and Kaplan, 2023; Vazquez et al., 2022), the equation of motion becomes

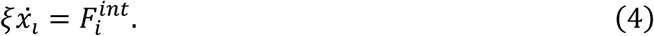

We assume the interaction force *F*^int^_i_ to be the sum of all pairwise interaction forces between cells and hence

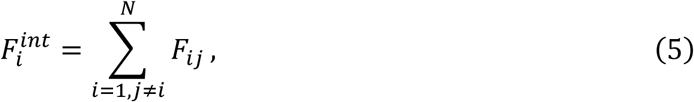

where *F*_ij_ is the interaction force between cells *i* and *j*. The pairwise interaction force is rotationally invariant and comprises both the attractive and repulsive components. An attractive component represents the adhesion between two cells, whereas a repulsive component opposes the deformation of cells owing to their elastic properties. Hence, we model the cell-cell interaction as a linear spring force together with a quadratic term that approximates the contact area between two cells (Delile et al., 2017) such that

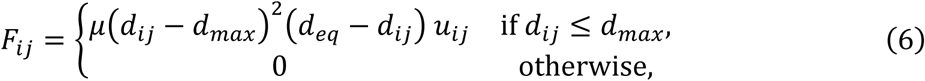

where μ is the cell interaction strength constant, *d*_ij_ is the distance between the two interacting cells defined as *d*_iJ_ = ‖*x*_i_ - *x*_J_‖, *d*_eq_ is the equilibrium distance defined as the sum of the radii of the two cells such that *d*_eq_ = *r*_i_ + *r*_J_, *d*_max_ is the maximum interacting distance between cells, hence defining the neighbors for each cell, and *u*_ij_ is the unit vector defining the force direction as 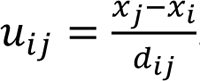.

### 2.2. Morphogens

Cells can express morphogens that diffuse into neighboring cells, which can dynamically change as cells move, divide, and die. The system of *N* cells can express *K* different morphogens, such as each cell *i* contains a concentration *m*_ik_ for each morphogen *k*, with *m*_ik_ ∈ ℝ. The concentration dynamics of each morphogen is defined as

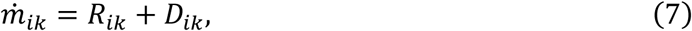

where *R*_ik_ is the reaction term and *D*_ik_ is the diffusion term of morphogen *k* in cell *i*. Intracellularly, morphogen *k* concentration in cell *i* changes due to production, degradation, and dilution, such as

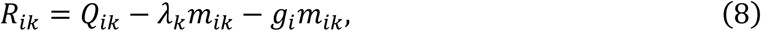

where *Q*_ik_ is the production term that depends on the particular regulation modeled, λ_k_ is the decay constant, and *g*_i_ is the growth rate of the cell. Morphogen diffusion occurs across neighboring cells and is approximated as a rate equation (Truskey et al., 2009), such as

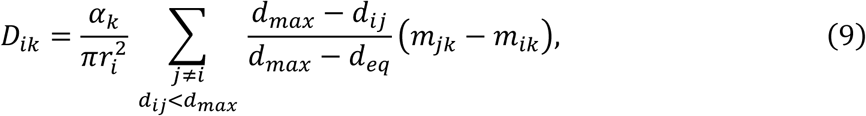

where α_k_ is the diffusion constant and π*r*^2^represents the cell area. In this way, the effective diffusion coefficient becomes α_k_ when two cells are at equilibrium distance *d*_eq_ from each other, but 0 when they are at or beyond their maximum interaction distance *d*_max_. The same cell center locations are the basis for calculating both the mechanical forces and morphogen diffusion fluxes, simplifying the mathematical complexity and computational requirements of the model.

### 2.3. Cell size dynamics

A growth morphogen with concentrations *m*_i1_ for each cell *i* regulates cell size dynamics. Depending on the local concentration of the growth morphogen, cells can grow or degrow exponentially, which in turn can trigger mitosis or apoptosis when their size is doubled or halved, respectively. Mitosis replaces the parent cell centered at *x*_i_ with two overlapping daughter cells, each half the parent size and centered at *x*_i_ ± (1 - √½) *r*_i_*u*_U_, where *u*_U_ is a unit vector with a uniform random direction, such that the daughter cells are maximally separated but within the area of the parent cell (Figure 1). Apoptosis removes the cell from the system. In addition, cell growth is modulated by local cell density, which can inhibit cell growth due to crowding. Thus, the growth rate *g*_i_ for cell *i* is defined as

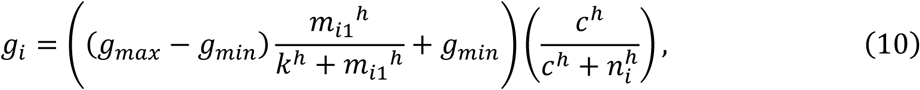

where *g*_max_ and *g*_min_ are the maximum and minimum cell growth rate constants, respectively,

*m*_i1_ is the growth morphogen concentration in cell *i*, *k* is the half-maximum concentration of the growth morphogen, *h* is the Hill coefficient, *n*_i_ is the number of neighbors of cell *i* within distance *d*_max_, and *c* is the half-maximum crowding inhibition. Then, the cell growth rate *g*_i_ modulates the area *A*_i_of cell *i* as

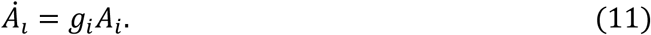

For all simulations, we set *g*_max_ = 0.01 and *g*_min_ = −0.01 so that the half-maximum concentration *k* denotes the growth morphogen concentration at which the growth rate is zero.

When the growth morphogen concentration is higher or lower than *k*, cells grow towards mitosis or shrink towards apoptosis, respectively. In addition, we set *h*=8.

**Figure 1.**
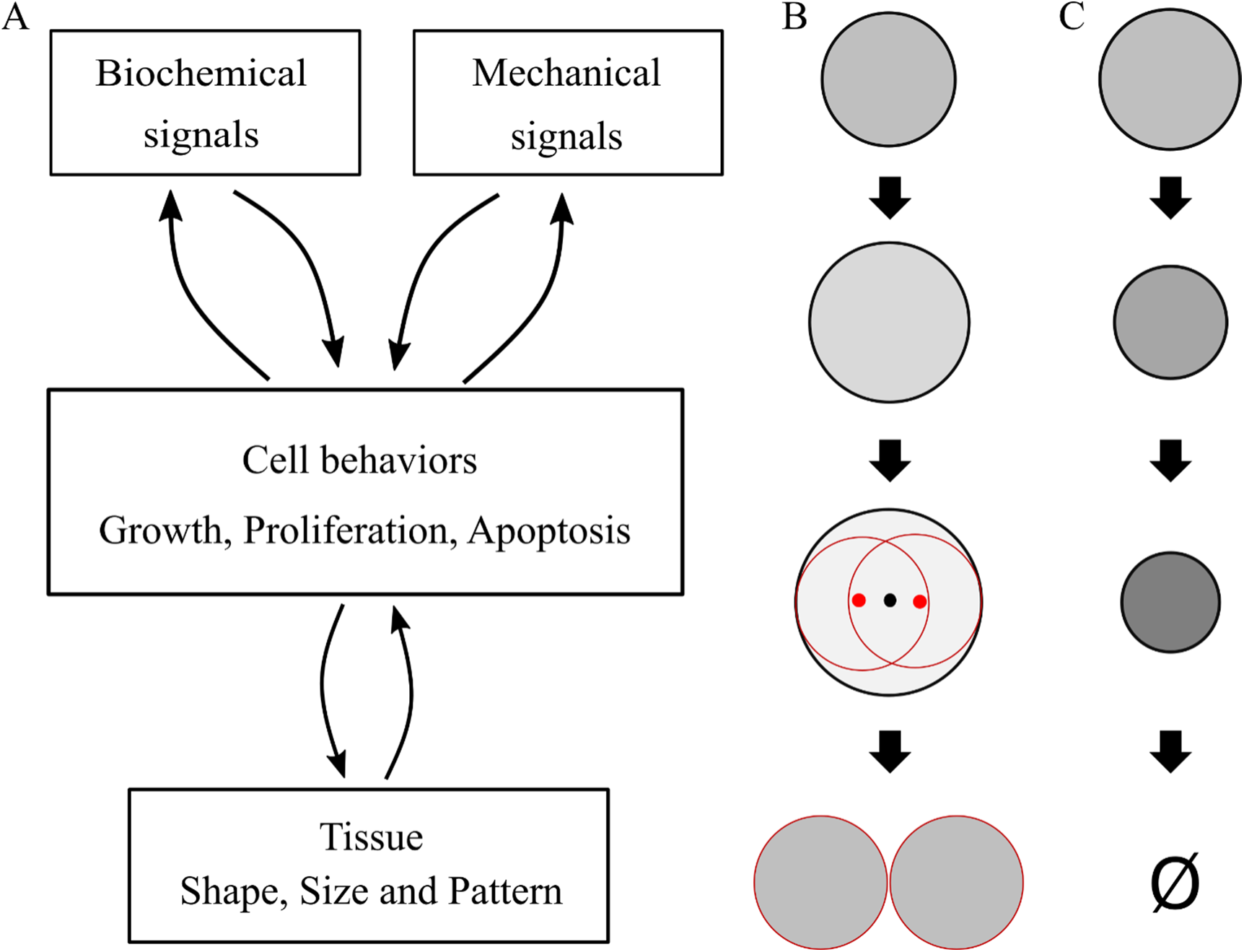
Proposed modeling framework for the regulation of whole-body shape dynamics. **A.** The model includes multiple layers of regulation that form feedback loops and give rise to emergent morphogen patterning and tissue morphogenesis. **B.** Cell area duplication results in mitosis. **C.** Cell area halving results in apoptosis.

### 2.4. Numerical methods

The system of equations was implemented and solved in MATLAB (MathWorks, Inc.) using the standard Euler forward scheme with a sufficiently small time step of Δ*t* = 0.001. To optimize the computational time, projection sorting (Law et al., 2019) was implemented to determine cell neighbors.

## 3. Results

### 3.1. Dynamic tissue shape modeling framework

To study the feedback regulation between morphogens and stable shapes in multicellular tissues and plastic whole organisms such as planarians, we present a modeling framework comprising off-lattice, agent-based cells that can migrate, grow, divide, and die as they are regulated by diffusive morphogens expressed by the cells (Figure 1). Cells interact with each other through attractive forces due to adhesion and repulsive forces due to deformation, which can produce stable tissue shape configurations. Crucially, morphogens can control cell growth rates and diffuse across cells, but not externally outside the tissue, which allows the modeling of self-contained systems with dynamic shapes, such as an organ or whole organism. In this way, the cells in the system can form one or more independent tissues, each made of a set of interconnected cells, where the morphogens can be expressed and diffuse within, but not across, the tissue boundary. In response to these morphogen concentrations, cells can grow or shrink, resulting in mitosis when the cell size is doubled or apoptosis when it is halved (Bortner and Cidlowski, 2002). The feedback loop between the biochemical signals forming spatial patterns and the mechanical forces between cells gives rise to the overall emergent shape of the whole tissue or organism, with the potential to reach a dynamic steady state when mitosis balances apoptosis.

### 3.2. Regulation of tissue shape and size by a single organizer cell

We sought to test the proposed model with a simulation of shape regulation from a single point source cell, an organizer, and investigate how tissue size and shape are modulated by different biological parameters. The initial state includes two cells: an organizer source cell expressing the diffusible growth morphogen at a constant rate *q* and a target cell (and its progeny) responding to the same morphogen. Hence, the production term *Q*_i1_ for each cell *i* is defined as

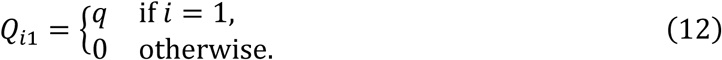

The morphogen expressed from the single source cell diffuses to neighboring cells and degrades at a constant rate (Equation 8). The source cell acts as an organizer with a constant volume, which prevents mitosis or apoptosis. The target cell and its progeny can divide or die through apoptosis in response to their size, which is regulated by the growth morphogen (Equation 10).

Figure 2A shows the tissue growth dynamic simulation with a single source cell (see Supplementary Video 1). The growth morphogen is expressed by the source cell (red) and diffuses through the cell population, creating a radial morphogen gradient. This gradient modulates the proliferative behavior of cells, resulting in tissue growth in all directions, which then surrounds the source cell at the center. The system reaches a steady state, forming a circular shape, in which the growth morphogen has a maximum concentration at the center, in the source cell, and a minimal concentration at the tissue shape edge. This concentration gradient produces a differential growth rate in the population, where cells near the center grow and divide, and those away from the center degrow and then die. At the steady state, the frequency of mitosis and apoptosis events balance out, and the number of cells in the tissue becomes constant. Thus, a single source cell causes the population to converge to a steady state with a circular shape.

**Figure 2.**
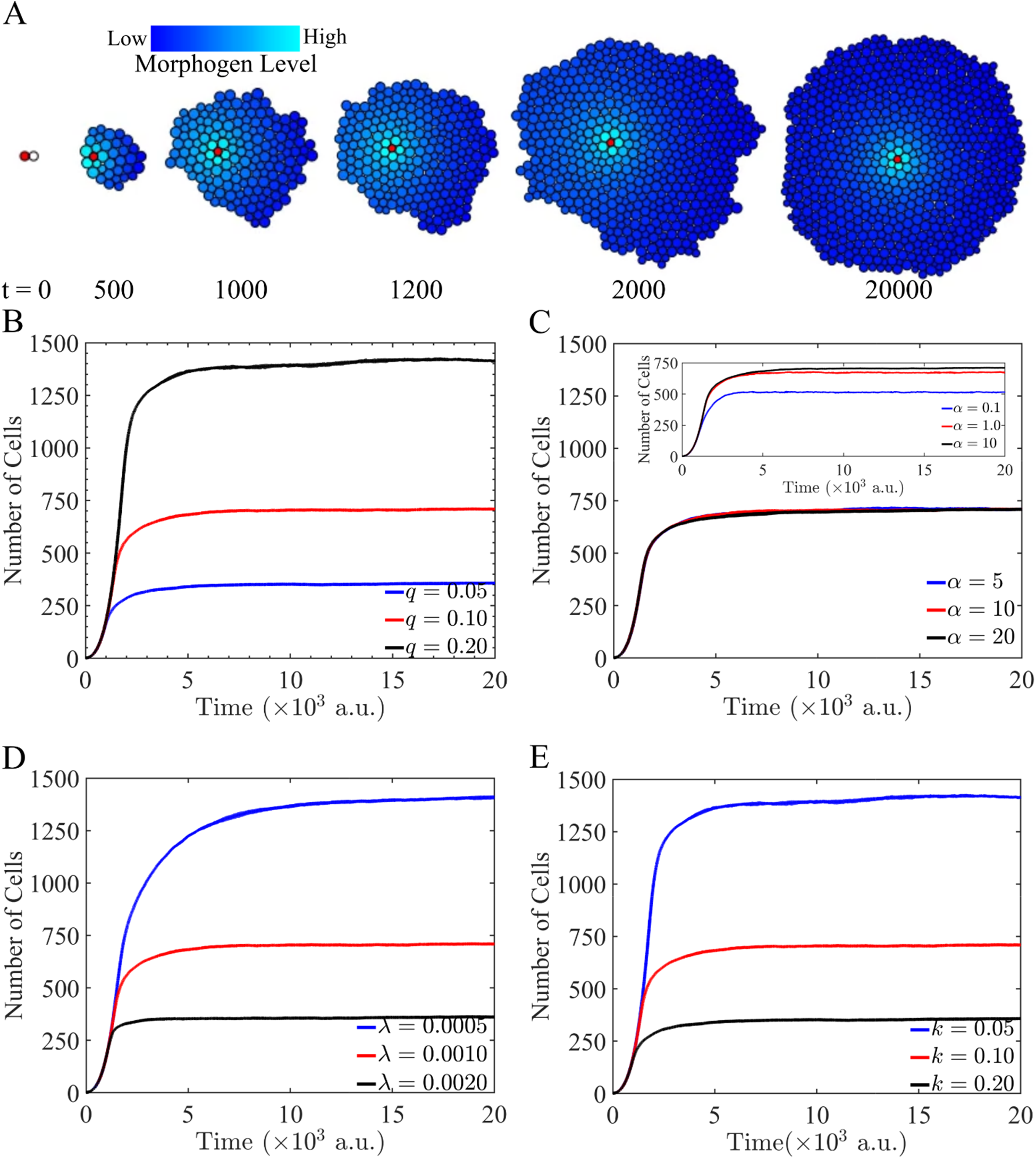
A single organizer source cell secreting a growth morphogen produces a steady-state circular shape that is modulated by the morphogen properties. **A.** Tissue growth controlled by a diffusing growth morphogen (blue) expressed from a single source cell (red) results in a steady-state circular shape. Parameters as red lines in panels B-E. **B.** The total number of cells at steady state depends linearly on the morphogen production constant (*q*). **C.** The morphogen diffusion constant (α) has a weaker effect on the total number of cells at the steady state. **D.** The morphogen degradation rate (λ) has an inverse effect compared to the production rate, albeit at a slower rate. **E.** The morphogen regulation threshold (*k*) between growth and degrowth has a similar but inverse effect compared to the morphogen production rate. All parameter sets were simulated ten times. The plots show the average (line) and standard deviation (shaded area). Except where indicated, the parameters were *q* = 0.1, α = 10, λ = 0.001, *R* = 0.1, and *c* = 6.

We next studied the effect of the morphogen dynamic parameters such as production, diffusion, degradation and regulation threshold on the cell and tissue emergent dynamics (Figure 2B-E). For each parameter set, ten simulations were performed—although the growth morphogen and cellular forces are deterministic, the cell division plane during mitosis is stochastic. All simulations resulted in a steady-state circular tissue shape, with the number of cells at steady state being modulated by the different biological parameters. The morphogen production constant linearly correlated with the number of cells (Figure 2B), whereas the morphogen degradation rate (Figure 2D) and regulation threshold (Figure 2E) were inversely correlated. Interestingly, the morphogen production constant and the regulation threshold had exactly opposite effects, since they both modulated the same regulatory dynamics (as modeled by a Hill function; see Equation 10). Although the morphogen diffusion constant was positively correlated with the number of cells (Figure 2C), its effect was much lower. In addition to the number of cells at steady state, the morphogen degradation rate also affected the speed at which the steady state was reached (Figure 2D), as a slower degradation rate results in slower equilibrium dynamics. All morphogen production and receptor parameters tested within a two-fold range resulted in a noticeable change in the number of cells at steady state. However, the morphogen diffusion coefficient only had a noticeable effect on the number of cells when it was changed by 10-fold (Figure 2C). Furthermore, as the diffusion coefficient modulates the morphogen diffusion length, a higher diffusion coefficient resulted in a larger diffusion area where cells grow and divide, causing an increased number of cells and tissue size at steady state (Figure 2C, inset). Crucially, there exists a critical value of the diffusion coefficient above which tissue growth dynamics remain unchanged, as the negative feedback between the growth morphogen concentration and tissue size is balanced.

In addition to morphogen parameters, the threshold of cell-crowding inhibition modulates the dynamics of shape formation. Figure 3 shows the simulations of tissue growth with a single-source morphogen at different cell-crowding inhibition thresholds. At a low inhibition threshold, growth was reduced at low local cell densities, which resulted in differential tissue expansion at the border towards the direction of low cell density, and hence, an irregular tissue shape (Figure 3A). In contrast, at high cell crowding thresholds, growth inhibition occurred only at high cell densities, resulting in more homogeneous growth across the tissue and faster convergence to a steady-state circular shape (Figure 3B). In addition to affecting shape dynamics, higher cell-crowding thresholds resulted in a larger number of cells at steady state, because cell growth was inhibited at higher densities (Figure 3C). Overall, the cell-crowding threshold plays a significant role in modulating tissue growth dynamics in terms of shape, size, and convergence speed (see Supplementary Video 2).

**Figure 3.**
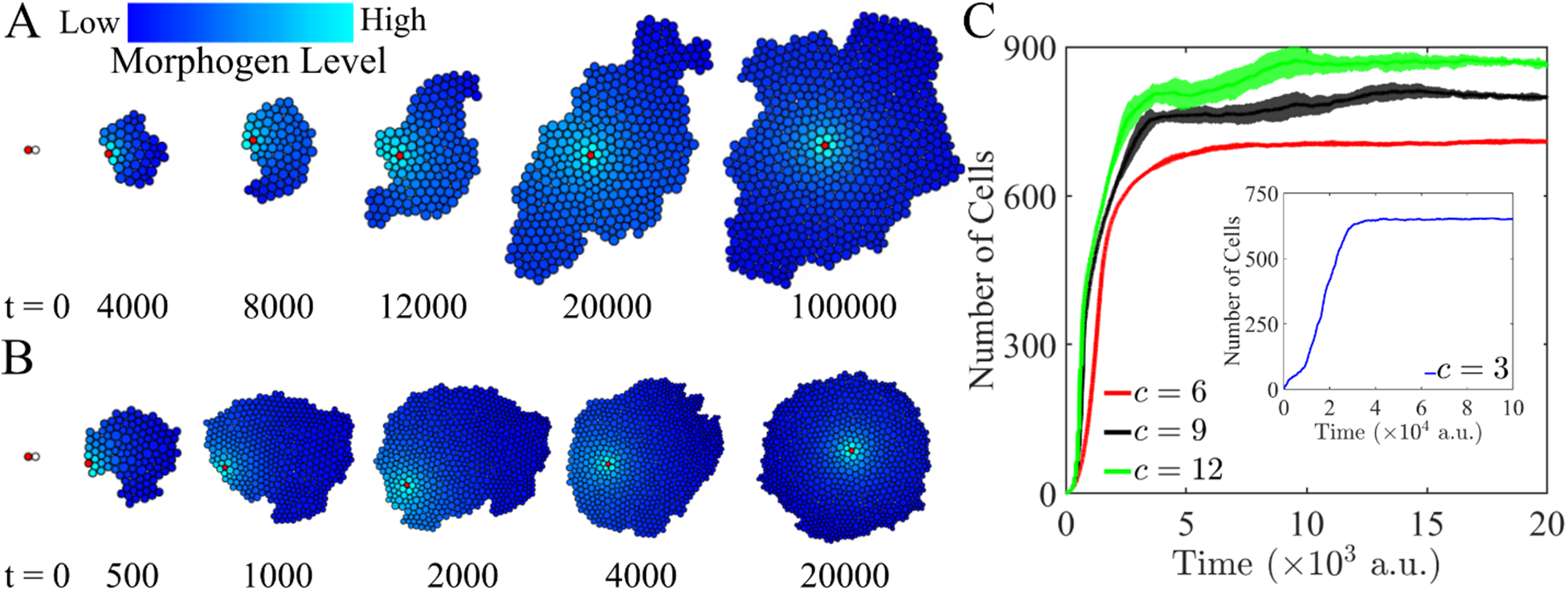
The shape dynamics regulated by a single organizer cell depend on the cell-crowding inhibition threshold. **A.** A low cell-crowding inhibition threshold (*c* = 3) inhibits growth at low local cell density, resulting in nonuniform cell proliferation that leads to irregular shapes. **B.** A high cell-crowding inhibition threshold (*c* = 12) inhibits growth only at a high local cell density, resulting in a circular shape. **C.** The growth curves show how higher cell-crowding inhibition thresholds result in a higher number of cells at steady state that are reached at shorter times. The inset shows the growth curve for a low cell-crowding inhibition threshold (*c* = 3), which requires a longer time to reach a steady state. All parameter sets were simulated ten times. The plots show the average (line) and standard deviation (shaded area). Remaining parameters: *q* = 0.1, α = 10, λ = 0.001, *R* = 0.1.

### 3.3. Regulation of tissue shape and size by multiple organizer cells

Plastic organisms such as planarians establish morphogen gradients secreted from multiple organizers, notably the anterior-posterior poles (Stückemann et al., 2017), which are essential for driving their whole-body shapes during homeostasis and regeneration (Gurley et al., 2008b; Ko et al., 2024; Reddien, 2018). Hence, we sought to determine how the number of organizer cells affects the steady-state shape of a tissue. Similar to the previous section, a single diffusible morphogen controls cell growth; however, more than one organizer expresses the growth morphogen. Figure 4 shows the simulations of tissue growth with two, three, or four organizer cells (red). As the growth morphogen concentration is higher near the source cells, higher proliferation rates occur in the cells surrounding the source cells. This higher proliferation rate, in turn, results in the separation of the source cells from each other until the growth morphogen diffuses to concentrations below its decay, which inhibits cell proliferation and promotes apoptosis. As the source cells acquire a stable distance from each other as the net cell proliferation and apoptosis balance, the resultant steady-state tissue shapes are not circular but can adopt elongated, triangular, or star shapes depending on the concentration pattern of the growth morphogen modulated by the number of source cells (see also Supplementary Video 3).

**Figure 4.**
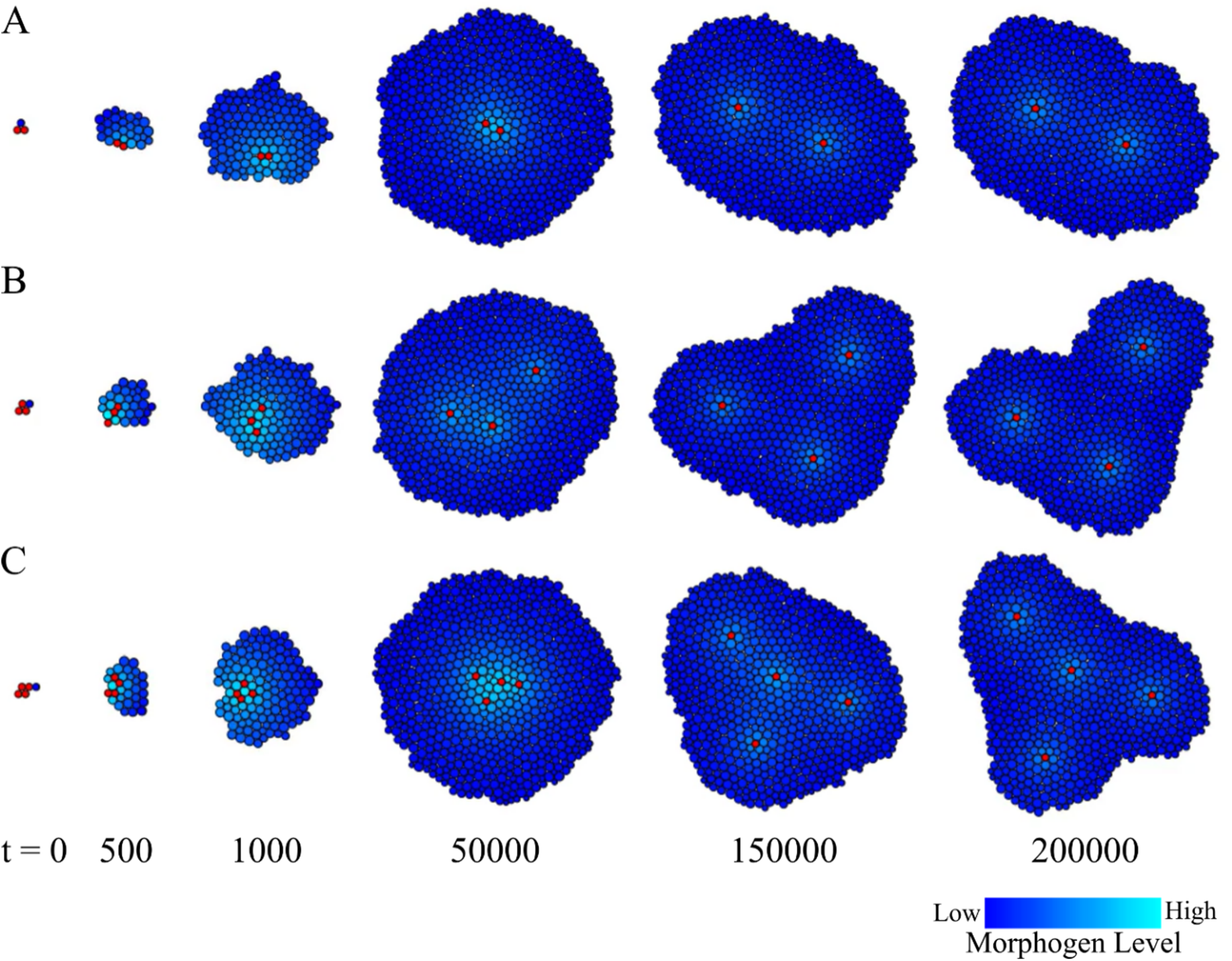
Multiple organizers secreting the same growth morphogen resulted in different steady-state tissue shapes. Cell proliferation induced by the secretion of the growth morphogen resulted in the organizers (red) moving away from each other. This repulsive effect modulates the concentration levels of the growth morphogen, resulting in oval (A), triangular (B), or star (C) steady-state tissue shapes under two, three, or four organizer cells, respectively. To maintain the number of cells at steady state equal, the morphogen production constant was scaled accordingly: *q* = 0.05, 0.033, or 0.025 for two, three, or four source cells, respectively. Remaining parameters: α = 10, λ = 0.001, *R* = 0.1, *c* = 6.

### 3.4. Formation of Turing spots and stripes patterns in a growing tissue

Morphogenesis and tissue patterning are not independent processes; rather, they act together through feedback loops to regulate gene expression, size, and shape in growing and differentiating tissues. Indeed, organizers need to be established during development and restored during regeneration, a process that is often dependent on self-organizing regulatory mechanisms, such as Turing systems patterning a dynamic tissue (Cotterell et al., 2015; Herath and Lobo, 2020; Maini et al., 2012; Marcon and Sharpe, 2012; Okuda et al., 2018b; Werner et al., 2015). We sought to shed light on the feedback process of self-regulation of tissue pattern, size, and shape using the proposed framework. First, we tested the capacity of the model to form spatial periodic patterns of morphogen expression with a regulatory network based on a self-regulated Turing mechanism. As before, a diffusing growth morphogen with concentrations *m*_i1_modulates cell growth dynamics and is expressed by a single source cell. Additionally, all cells express two other morphogens with concentrations *m*_i2_and *m*_i3_, respectively, that interact and diffuse across neighboring cells based on Schnakenberg kinetics (Schnakenberg, 1979). Continuous reaction-diffusion systems modeled using Schnakenberg kinetics can produce spot or stripe patterns over a relatively large Turing parameter space compared to other Turing models. In this system, the first morphogen self-promotes its expression while inhibiting the expression of the second morphogen. Conversely, the second morphogen inhibits its own expression and promotes the expression of the first morphogen. In this way the production terms of the Turing morphogens are defined as

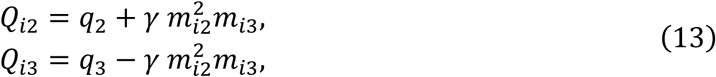

where *q*_2_ and *q*_3_ are the basal expression levels of each morphogen, and γ is a coupling constant. In addition, we introduce a scaling parameter Λ in the Turing morphogens’ reaction term that can control the spatial scale of the periodic pattern, such as

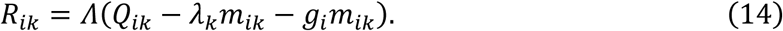

Figure 5 shows the simulations of the spatial patterning dynamics over time in a proliferating and growing tissue domain from a single source cell for the growth morphogen (see Supplementary Video 4). The growth morphogen from a single organizer cell modulates tissue growth dynamics, resulting in a stable circular shape. As all cells implement the reaction-diffusion Turing mechanism, the proliferative tissue dynamically forms a spot pattern that, concomitant with the tissue shape, reaches a stable steady state. The wavelength of the system, that is, the distance between the spot centers, is modulated by the scaling parameter Λ, which also influences the spot size. A small scaling factor Λ = 0.1 results in a higher wavelength, leading to a single stable spot at the center of the tissue domain (Figure 5A). As the scaling factor Λ is increased to 0.5, the wavelength and spot size decrease, resulting in three stable spots in the same tissue domain (Figure 5B). A further increase in the scaling factor Λ to 2.0 results in a higher number of smaller spots (Figure 5C).

**Figure 5.**
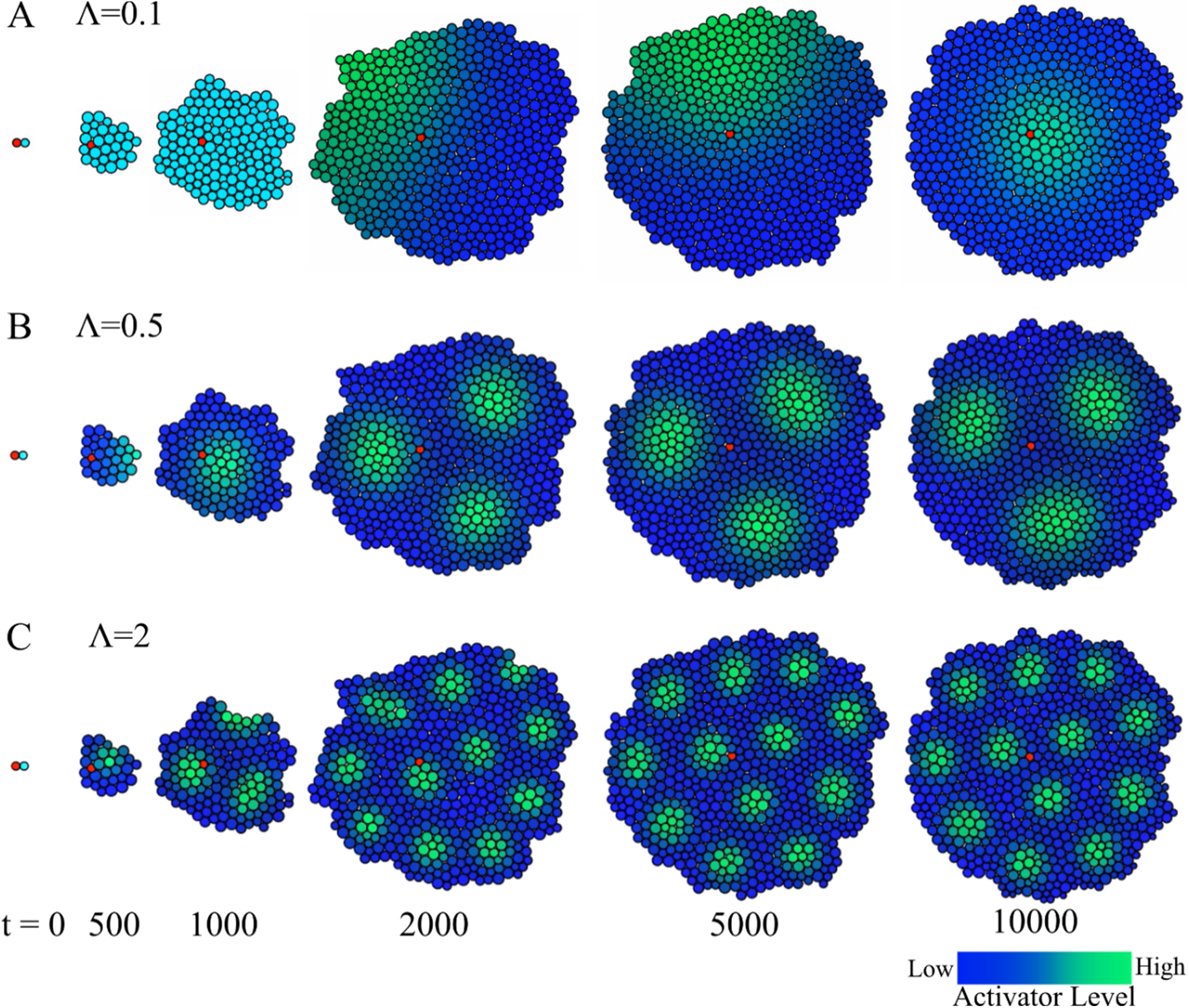
A growing tissue with a single organizer can produce stable Turing spot patterns at different wavelengths. A single organizer cell (red) secretes a diffusive growth morphogen, resulting in a stable circular tissue shape. In addition, all cells include a genetic circuit implementing an activator-inhibitor Schnakenberg Turing mechanism that reacts and diffuses across the domain. The system can produce spot patterns across the growing tissue, reaching a stable steady-state spatial configuration. **(A-C)** The scaling factor Λ modulates the number and size of the spots. Blue-green indicates the concentration of the activator. Parameters: *q*_1_ = 0.1, *q*_2_ = 0.1, *q*_3_ = 0.9, α_1_ = 10, α_2_ = 2, α_3_ = 40, λ_1_ = 0.001, λ_2_ = 1.0, λ_3_ = 0.1, γ = 1.0.

In addition to spot patterns, a Turing mechanism can produce stripes in a continuous domain. To test the ability of the proposed model to produce stable stripe patterns in a growing tissue, the production term of the inhibitor morphogen was modified such that it is modulated by the activator morphogen, resulting in

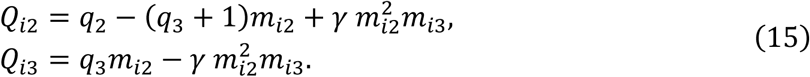

Figure 6 shows the simulations of the modified Schnakenberg system over time in a proliferating and growing tissue domain from a single organizer cell secreting the growth morphogen (see Supplementary Video 5). As with the spot pattern, the growth morphogen signaling resulted in a stable circular shape, whereas the reaction-diffusion mechanism dynamically produced a stable stripe pattern. Similarly, the scaling parameter Λ modulates the wavelength of the system—in this case, the distance between the stripes. Hence, at the steady state, increasing the values of the scaling parameter Λ resulted in one, three, or multiple stripes (Figure 6A-C, respectively). Remarkably, the spot and stripe patterns did not form at high scaling parameter values (data not shown) because the system wavelength becomes smaller than the size of a single cell. Conversely, low values of the scaling parameter may require domain sizes larger than the tissue size; otherwise, the spot and stripe patterns do not form. These analyses demonstrate the versatility of the model for studying periodic pattern formation in dynamic tissue shapes.

**Figure 6.**
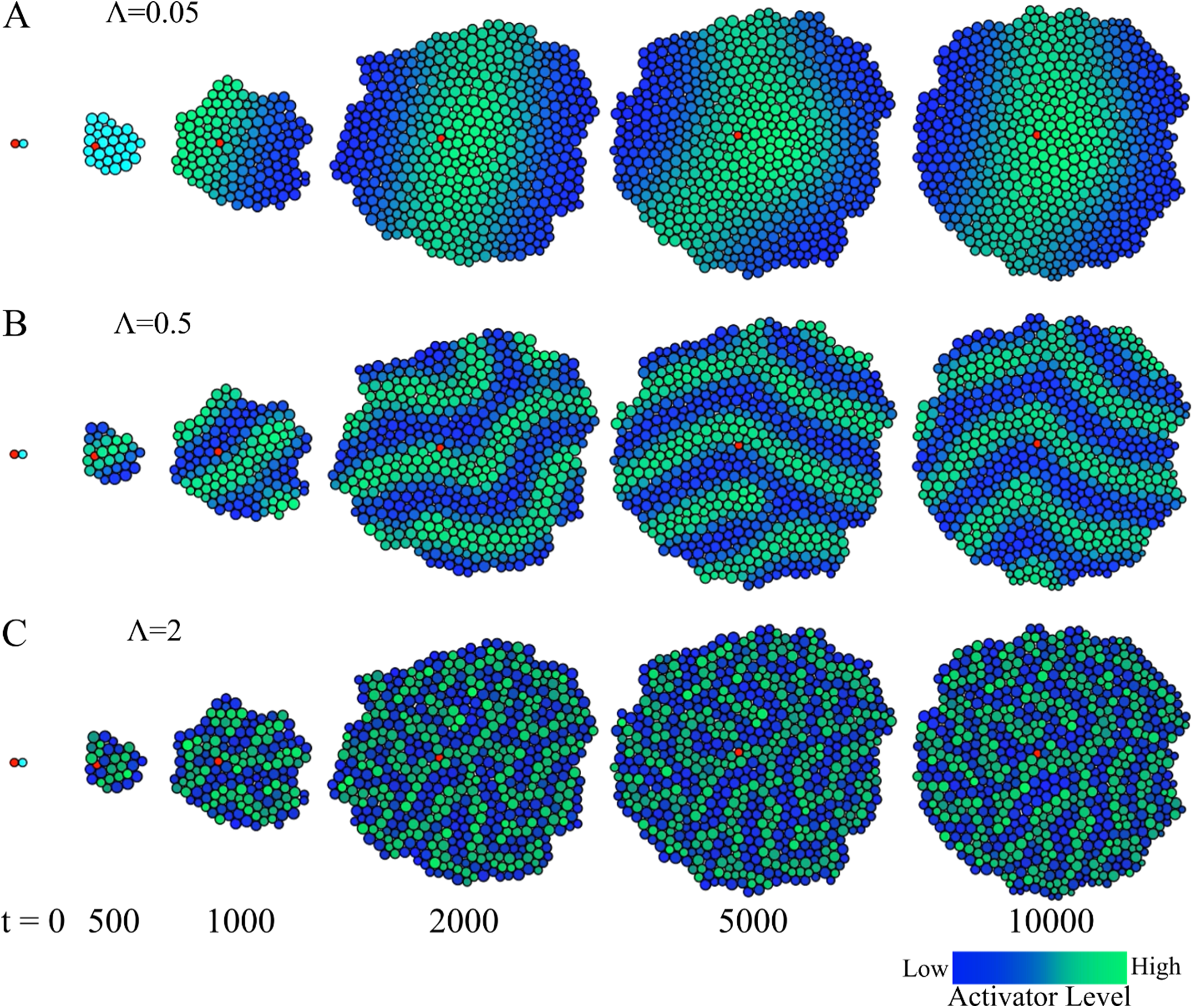
Stable Turing stripe patterns at different wavelengths can also be formed in growing tissues with a single organizer cell. When the production term of the inhibitor is also modulated by the activator in the Schnakenberg Turing mechanism, the growing tissue with a single organizer cell (red) can produce stable stripe patterns instead of spot patterns (see Figure 5). **(A-C)** The scaling factor Λ modulates the number and size of the stripes. Blue-green indicates activator concentration. Parameters: *q*_1_ = 0.1, *q*_2_ = 3, *q*_3_ = 9, α_1_ = 10, α_2_ = 3, α_3_ = 10, λ_1_ = 0.001, λ_2_ = 0, λ_3_ = 0, γ = 1.0.

### 3.5. Tissue shape controlled by Turing mechanisms

The coupling of gene expression patterning and growth to regulate the shape of a tissue or whole body is a complex process that is not well understood. Genes such as sonic hedgehog can act as both morphogens and mitogens to regulate vertebrate development, including patterning, proliferation, growth, and differentiation (Groves et al., 2020). In planarians, positional control genes form morphogen gradients across the three body axes (Bonar et al., 2022; Reddien, 2021; Witchley et al., 2013). Notably, incorrect morphologies and whole-body shapes can result from perturbations in the Wnt/β-catenin pathway (Gurley et al., 2008a; Scimone et al., 2016; Sureda-Gómez et al., 2016), which has been hypothesized to be established with a Turing mechanism (Herath and Lobo, 2020; Stückemann et al., 2017; Werner et al., 2015). Hence, modeling the complex interplay between pattern formation, cell proliferation, and tissue growth can provide mechanistic insights into these feedback regulatory processes (Economou et al., 2012; Krause et al., 2023). To investigate the emergence of shapes through shape-morphogen feedback mechanisms, we adapted the proposed Turing system to produce spatial patterns that act as morphogens that control cell growth. In this way, no predetermined organizer cells express the growth morphogen; instead, all cells are regulated by the same activator-inhibitor system (Equation 13), whereas the activator acts as the growth morphogen and directly regulates cellular growth (Equation 10).

Figure 7 shows different simulations of the Turing system regulating cellular growth dynamics, all resulting in different stable tissue shapes depending on the duration of the developmental period (see also Supplementary Video 6). The initial state included a single cell with equal activator-inhibitor system concentrations. A low growth morphogen threshold *R* = 0.1 (Equation 10) induces cell growth and proliferation during development. As the tissue grows, the Turing system produces different spot patterns that, in turn, regulate cellular proliferation in a feedback loop. After the developmental period, the growth morphogen threshold increases to *R* = 0.4, which reduces cell proliferation and leads to a steady-state Turing pattern and tissue shape. Figure 7A shows how a short developmental period results in a circular shape with a single Turing morphogen spot that regulates cell proliferation. The high concentration of this growth morphogen at the tissue center (green cells) results in a higher cell proliferation rate that balances the higher apoptosis rate at the border (blue cells) due to the lower growth morphogen concentration. The proliferation at the center balances with the apoptosis at the border and results in a dynamically stable steady state, forming a circular tissue shape. Hence, the circular stable spatial tissue configuration is driven by a feedback loop between the Turing pattern that controls cell proliferation and the resulting tissue shape, where the Turing pattern reacts and diffuses.

**Figure 7.**
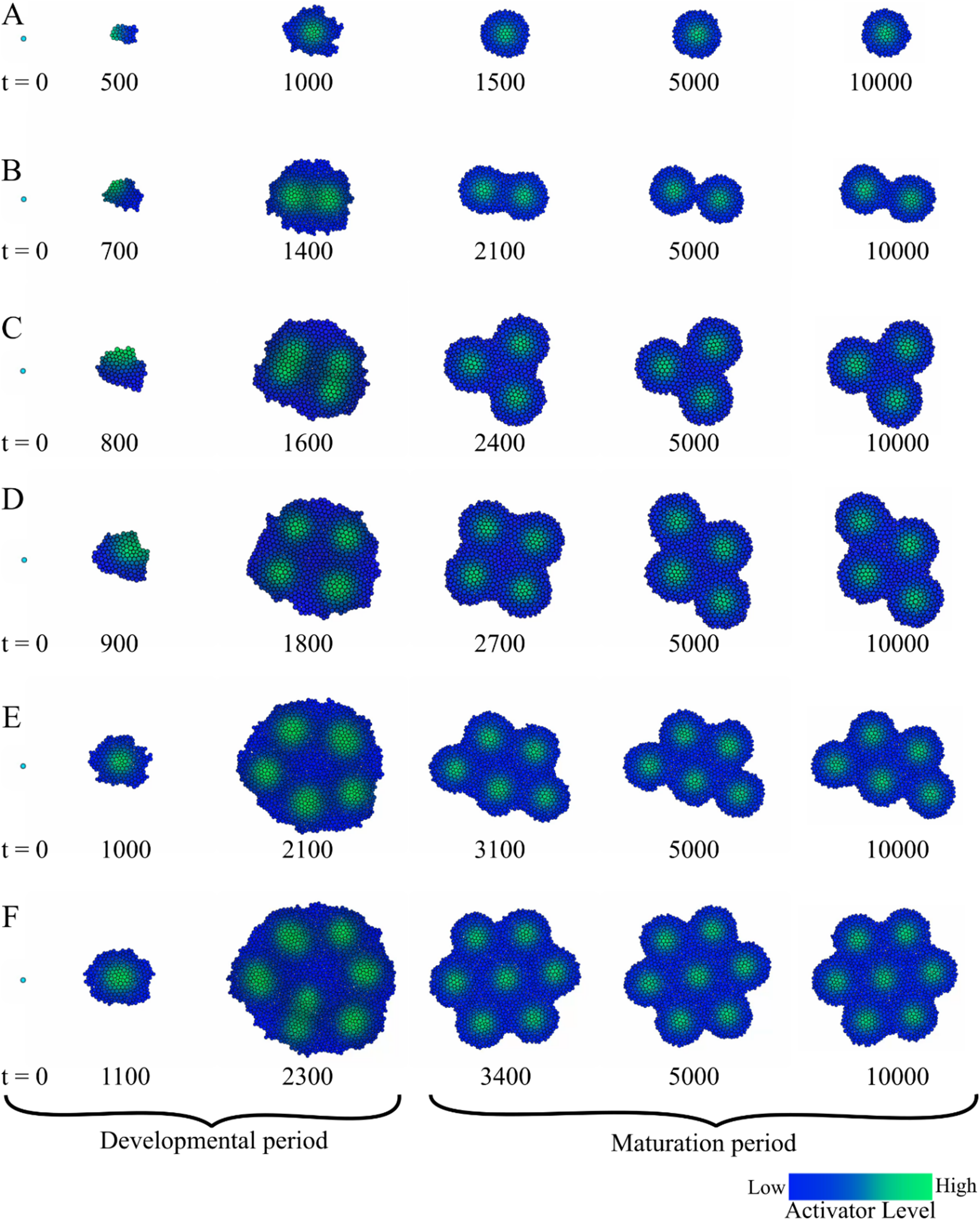
The feedback between a Turing spot mechanism controlling cell proliferation and the resulting tissue growth can produce different stable shapes depending on the length of the developmental period. (A-F) Simulation snapshots of tissue growth governed by Schnakenberg activator-inhibitor kinetics. The Turing system drives both the spot spatial pattern and the resulting tissue shape, as the activator morphogen regulates cell growth. A first period of development with high cell proliferation (growth morphogen threshold *R* = 0.1) causes an initial tissue growth, after which a stable dynamic shape is established owing to a lower cell proliferation (*R* = 0.4). The length of the developmental period dictates the number of spots developed and hence the final tissue shape. Parameters: *q*_2_ = 0.1, *q*_3_ = 0.9, α_2_ = 2, α_3_ = 40, λ_2_ = 1.0, λ_3_ = 0.1, γ = 1.0.

Crucially, the length of the developmental period modulates the number of spots, and hence the resulting final shape at the steady state (Figure 7 B-F). Longer developmental periods increase proliferation, which results in a higher number of cells at the stable steady state. Because the wavelength of the Turing system is constant, a larger number of spots develop in larger tissues, which in turn can produce different shapes. Figure 7 shows simulations with increased developmental periods that led to stable steady-state tissue patterns and shapes with one to seven spots. The shapes resulting from the different numbers of spots included circular, hourglass, triangular, and up to hexagonal for one, two, three, and seven spots, respectively.

In addition to spot patterns, the Schnakenberg system can produce stripe patterns (Figure 6), which can act as morphogen signals for cell proliferation. Figure 8 shows simulations of the stripe-producing Schnakenberg system, where the activator regulates cell growth (see also Supplementary Video 7). As before, the initial state included a single cell with equal morphogen concentrations. The simulation was divided into a developmental period with a low morphogen threshold (*R* = 0.5), resulting in high cell proliferation, followed by a stable period with a high morphogen threshold (*R* = 2.5), causing a balance between cell proliferation and apoptosis. Although the number of cells reached a steady state, the stripe pattern, in contrast to the spot pattern, did not result in a stable tissue shape. Because the activator stripes reach the edge of the tissue, cell proliferation is high at these locations, leading to shape instability and dynamic continuous reconfiguration of the stripe pattern. The overall size of the tissue is controlled by the length of the developmental period; however, the resulting dynamic tissue shapes are similar.

**Figure 8.**
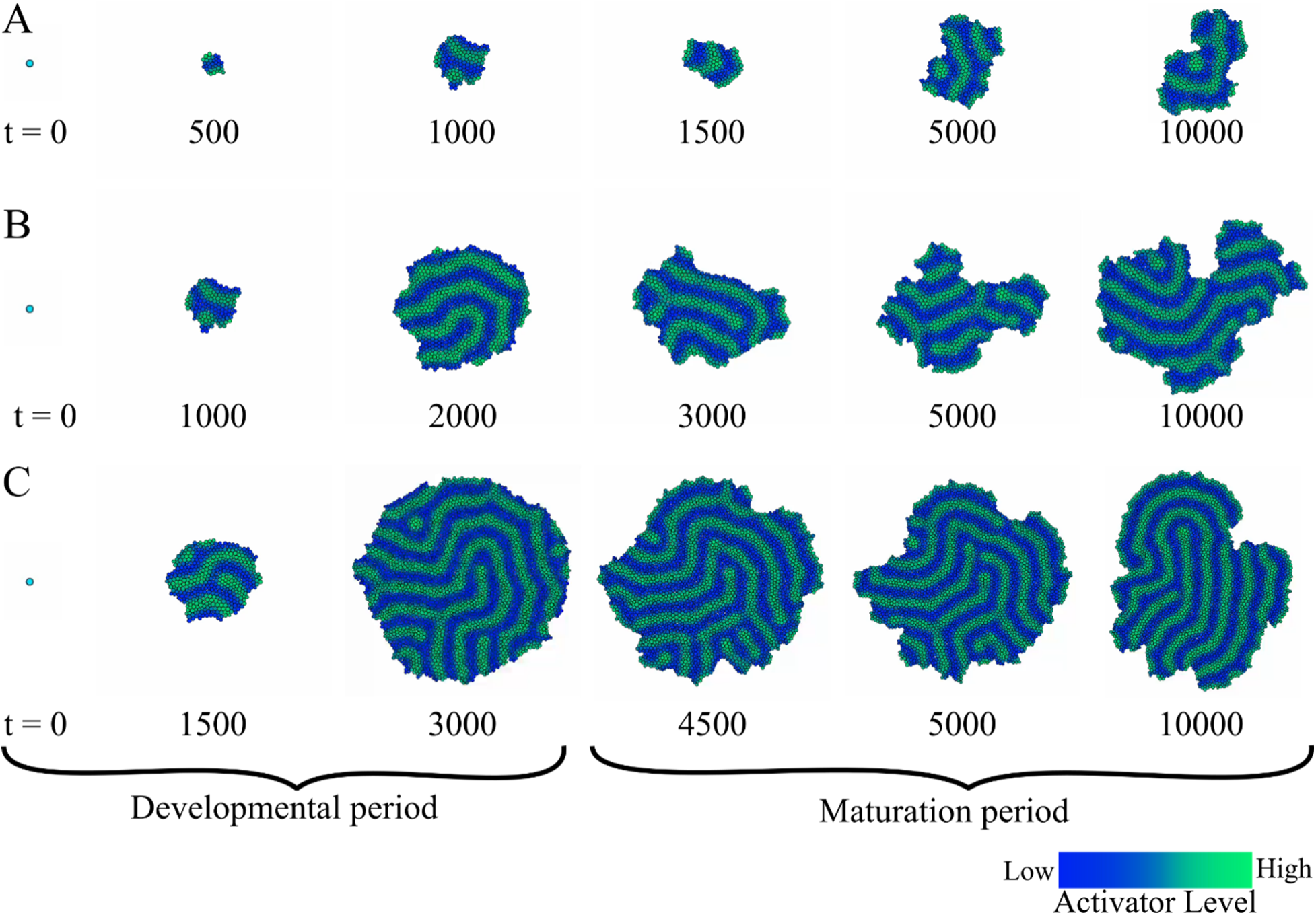
A Schnakenberg stripe mechanism controlling cell proliferation results in dynamic tissue shapes. (A-C) Simulation snapshots of tissue growth governed by Schnakenberg activator-inhibitor kinetics driving the stripe spatial pattern and resulting tissue shape. During the developmental period (growth morphogen threshold *R* = 0.5), the tissue grows, followed by a stable period (*R* = 0.4) when cell proliferation and apoptosis are balanced. An increased developmental period results in larger tissue sizes; however, the tissue shape dynamically changes owing to stripe-driven cell proliferation at the tissue border. Parameters: *q*_2_ = 3, *q*_3_ = 9, α_2_ = 3, α_3_ = 10, λ_2_ = 0, λ_3_ = 0, γ = 1.0.

Although the stripe-forming Schnakenberg mechanism, which controls cell proliferation, produces globular tissue shapes (Figure 8), the stripe Turing system has the potential to produce a single elongated shape. For this, a single stripe with a high activator concentration at the midline and a low concentration at the border could form a stable elongated shape. Hence, the high proliferation rate at the midline could balance apoptosis at the border to form a stable filament shape. At the extreme of the stripes, a high concentration of the activator could induce stripe growth. We tested this hypothesis with a molecularly plausible Gierer-Meinhardt model of reaction-diffusion mechanisms (Gierer and Meinhardt, 1972; Koch and Meinhardt, 1994; Yamaguchi et al., 2007) defined as

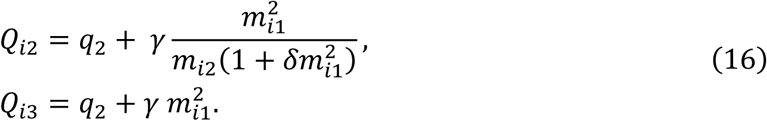

where *q*_1_ and *q*_2_ are the basal expression levels of each morphogen, γ is a coupling constant, and δ is the inverse of the activator production threshold. The scaling parameter Λ controls the spatial scale of the periodic pattern (Equation 14).

Figure 9 shows the simulations of the Gierer-Meinhardt model, where the activator morphogen regulates cell growth (see also Supplementary Video 8). Starting with a single cell, the tissue proliferates isometrically until a stripe pattern is formed. The high proliferation rate at the stripe midline was balanced with the high apoptosis rate at the lateral border, resulting in a filament shape (Figure 9B). The activator inducing a high proliferation rate is also high at the tip of the filament, which causes continuous growth of the filament. As the scaling factor Λ decreased to 0.05, both the wavelength and stripe size increased, resulting in wider stripes within the tissue domain (Figure 9A). Conversely, increasing the scaling factor Λ to 0.5 leads to a higher number of thinner stripes, resulting in a more circular shape (Figure 9C).

**Figure 9.**
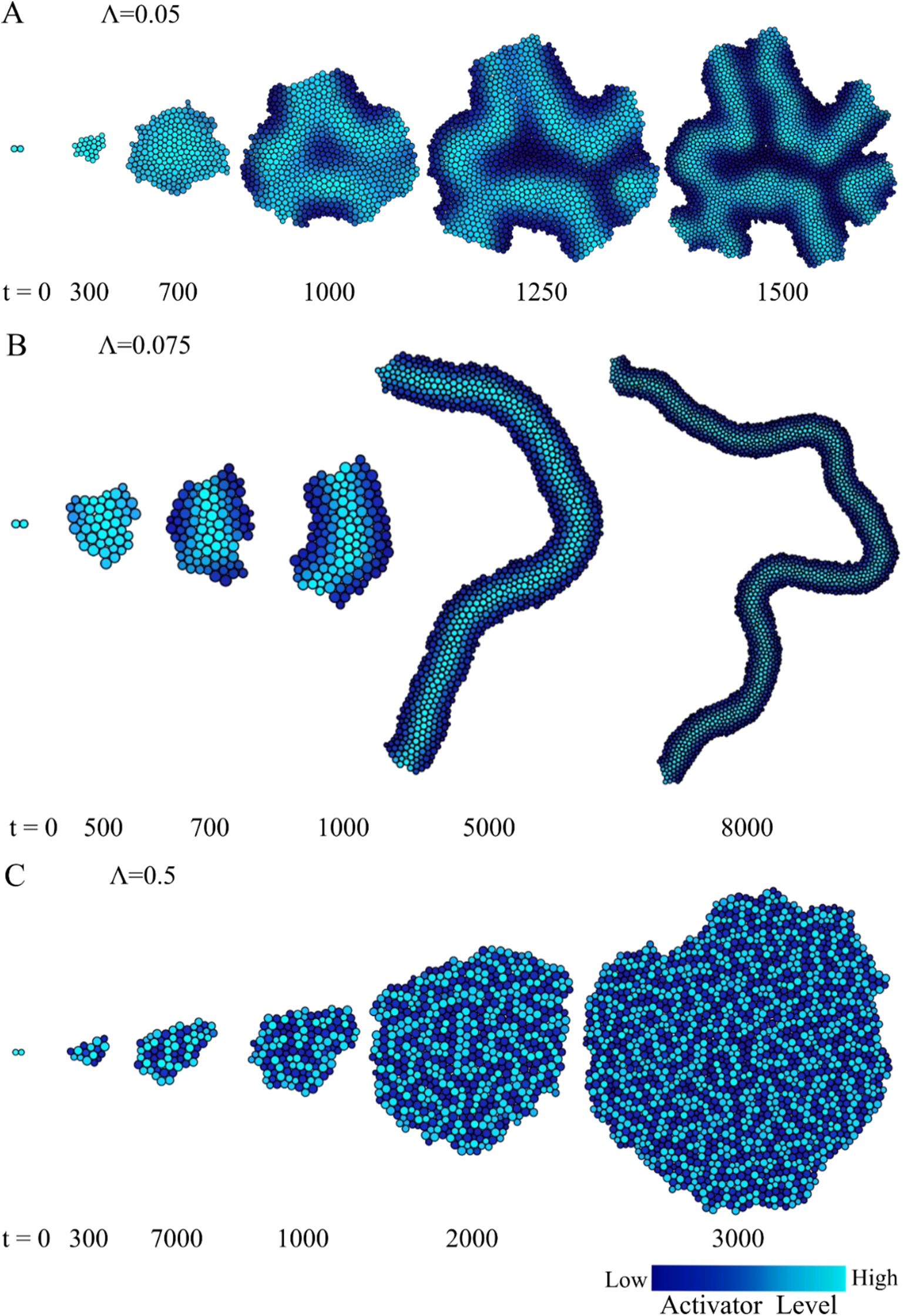
A Gierer-Meinhardt stripe mechanism controlling cell proliferation resulted in growing filament tissue shapes. Simulation snapshots of tissue growth governed by Gierer-Meinhardt reaction-diffusion kinetics driving the formation of a filament tissue shape formed by a single stripe. **(A-C)** The scaling factor Λ modulates the pattern and shape. Parameters: *q*_2_ = 0, *q*_3_ = 0, α_2_ = 0.1, α_3_ = 2, λ_2_ = 1.2, λ_3_ = 1.0, γ = 1.0, δ = 0.4.

## 4. Discussion

Here, we investigated the role of the feedback loop between mechanochemical signaling pathways and cellular growth to regulate the formation of stable tissue shapes. We focused on how regulatory signals that diffuse and react in the tissue can drive the formation of steady-state emergent shapes and patterns through the balance of mitosis, apoptosis, and cellular forces, as observed during the development of organs or whole-body shape dynamics of plastic organisms such as planarians. A mathematical framework for morphogen-mediated tissue growth dynamics was developed to demonstrate how cell-level phenomena, including growth, proliferation, and apoptosis, influence tissue-scale dynamics. In turn, the overall tissue shape and size feedback into the regulation of morphogen patterns as they react and diffuse through the tissue formed by the cells, closing the loop between cellular behaviors and emergent shapes. As the model uses the cell-center positions for calculating both mechanical forces and chemical diffusion, it allows for the efficient simulation of independent tissues, such as organs or whole bodies, driven by morphogens forming spatial patterns within the tissue. The model also includes cell death and proliferation, which are essential for the formation of steady-state tissue configurations in terms of shape, size, and patterning.

The proposed framework was employed to study the biological drivers that regulate the feedback loop between morphogens and tissue shapes through cell growth. The results showed that different tissue shapes emerged depending on the number of source cells (organizers) that secrete diffusive morphogens. For a single source cell, a stable circular tissue shape was formed at the steady state, whereas a stable elliptical tissue shape reminiscent of elongated body shapes, such as planarians, resulted from two source cells. As the number of source cells increased, the shape became increasingly complex. Parametric analysis of the single-source cell system revealed that the morphogen diffusion constant, degradation rate, and regulation threshold play crucial roles in tissue growth dynamics and final shape size. Finally, the results showed that the time to reach a steady state and hence a stable tissue size are highly sensitive to the parameters that regulate intracellular morphogen dynamics and their microenvironment.

As biological systems can produce and restore shapes and patterns during homeostasis and regeneration, the model was then extended to investigate self-regulated feedback mechanisms between patterning and cellular behaviors, and their role in the regulation of stable tissue shapes. First, we studied how the dynamics of a Turing activator-inhibitor system can evolve concomitantly with tissue growth to produce stable patterns. The results showed how cell fate specification can emerge from an intracellular activator-inhibitor system as the pattern forms and scales with the overall tissue size. Both spot and stripe patterns were observed, depending on the regulatory interactions between morphogens in the Turing system. Crucially, when the feedback loop between cell behavior and tissue patterning was closed by replacing the organizer cells with a self-organized Turing morphogen regulating cell growth, different stable tissue shapes could be formed. In the spot pattern dynamics, various stable morphological states, from circular to elongated to cluster-like, were observed depending on the length of the developmental period. In contrast, the stripe pattern dynamics resulted in dynamic tissue shapes, including growing filaments, owing to the presence of a growing front at the tissue border. These results provide insights into the coupling between mechanochemical signals and tissue dynamics to form self-regulated stable tissue shapes by spatially balancing mitosis and apoptosis with a Turing system.

This study focused on tissue growth and stable shape formation by balancing cell proliferation and apoptosis. Such a balance of cell numbers allows plastic organisms such as planarians to grow and degrow their adult bodies by more than an order of magnitude while maintaining correct body shapes (Baguñà et al., 1990; Hill and Petersen, 2015; Ko et al., 2024; Stückemann et al., 2017). Although gradient-forming morphogens are essential for correctly patterning the planarian body (Reddien, 2021), how such patterns form and feedback into the tissue growth and shape they control remains poorly understood. This study sheds light on possible feedback mechanisms between morphogens and tissue growth to form such self-regulated shapes. Future work will extend the presented approach with specific regulatory networks and quantitative phenotypes towards a mechanistic understanding of whole-body shape and scaling regulation. Crucially, the proposed approach can be integrated into machine learning algorithms that can infer models for particular patterns and shapes (Mousavi et al., 2021; Mousavi and Lobo, 2024). These predictive models can then be validated experimentally at the level of whole-body tissue phenotypes (Lobo, 2022; Roy et al., 2020) and gene expression patterns using whole-mount in situ hybridization assays (Wolff et al., 2022). We have limited this theoretical study to two-dimensional tissue shapes and patterns, which is ideal for the anterior-posterior and medio-lateral phenotypes mostly available in organisms such as planarians. However, the proposed framework can be easily extendable to three-dimensional studies of self-regulated stable morphogenesis.

Overall, the analysis presented in this study provides evidence of the significant role of the regulatory feedback between morphogens and cell dynamics in controlling target tissue and whole-body shapes during morphogenesis. The proposed framework linking cellular behaviors, patterning, and tissue shapes provides a robust theoretical basis for studying dynamic tissue morphogenesis and patterning.

## Acknowledgements

We thank the members of the Lobo Lab for helpful discussions. This work was supported by the National Institute of General Medical Sciences of the National Institutes of Health under award number R35GM137953. The content is solely the responsibility of the authors and does not necessarily represent the official views of the National Institutes of Health. Computations used the UMBC High Performance Computing Facility (HPCF) supported by the NSF MRI program grants CNS-1920079 and OAC-1726023.

## References

Adell, T., Cebrià, F., Abril, J.F., Araújo, S.J., Corominas, M., Morey, M., Serras, F., González-Estévez, C., 2025. Cell death in regeneration and cell turnover: Lessons from planarians and *Drosophila*. Seminars in Cell & Developmental Biology 169, 103605. 10.1016/j.semcdb.2025.103605

Aegerter-Wilmsen, T., Heimlicher, M.B., Smith, A.C., De Reuille, P.B., Smith, R.S., Aegerter, C.M., Basler, K., 2012. Integrating force-sensing and signaling pathways in a model for the regulation of wing imaginal disc size. Development 139, 3221–3231. 10.1242/dev.082800

Baguñà, J., Romero, R., Saló, E., Collet, J., Auladell, C., Ribas, M., Riutort, M., García-Fernàndez, J., Burgaya, F., Bueno, D., 1990. Growth, Degrowth and Regeneration as Developmental Phenomena in Adult Freshwater Planarians, in: Experimental Embryology in Aquatic Plants and Animals. Springer US, Boston, MA, pp. 129–162. 10.1007/978-1-4615-3830-1_7

Bailles, A., Gehrels, E.W., Lecuit, T., 2022. Mechanochemical Principles of Spatial and Temporal Patterns in Cells and Tissues. Annual Review of Cell and Developmental Biology 38, 321–347. 10.1146/annurev-cellbio-120420-095337

Bonar, N.A., Gittin, D.I., Petersen, C.P., 2022. Src acts with WNT/FGFRL signaling to pattern the planarian anteroposterior axis. Development 149, dev200125. 10.1242/dev.200125

Bortner, C.D., Cidlowski, J.A., 2002. Apoptotic volume decrease and the incredible shrinking cell. Cell Death & Differentiation 9, 1307–1310. 10.1038/sj.cdd.4401126

Boulet, A.M., Moon, A.M., Arenkiel, B.R., Capecchi, M.R., 2004. The roles of Fgf4 and Fgf8 in limb bud initiation and outgrowth. Developmental Biology 273, 361–372. 10.1016/j.ydbio.2004.06.012

Cano-Fernández, H., Tissot, T., Brun-Usan, M., Salazar-Ciudad, I., 2025. A mathematical model of development shows that cell division, short-range signaling and self-activating gene networks increase developmental noise while long-range signaling and epithelial stiffness reduce it. Developmental Biology 518, 85–97. 10.1016/j.ydbio.2024.11.014

Cockerell, A., Wright, L., Dattani, A., Guo, G., Smith, A., Tsaneva-Atanasova, K., Richards, D.M., 2023. Biophysical models of early mammalian embryogenesis. Stem Cell Reports 18, 26–46. 10.1016/j.stemcr.2022.11.021

Collinet, C., Lecuit, T., 2021. Programmed and self-organized flow of information during morphogenesis. Nat Rev Mol Cell Biol 22, 245–265. 10.1038/s41580-020-00318-6

Cotterell, J., Robert-Moreno, A., Sharpe, J., 2015. A Local, Self-Organizing Reaction-Diffusion Model Can Explain Somite Patterning in Embryos. Cell Systems 1, 257–269. 10.1016/j.cels.2015.10.002

Davies, E.L., Lei, K., Seidel, C.W., Kroesen, A.E., McKinney, S.A., Guo, L., Robb, S.M.C.M.M.C., Ross, E.J., Gotting, K., Alvarado, A.S., Sánchez Alvarado, A., 2017. Embryonic origin of adult stem cells required for tissue homeostasis and regeneration. eLife 6, 1–35. 10.7554/eLife.21052

Delile, J., Herrmann, M., Peyriéras, N., Doursat, R., 2017. A cell-based computational model of early embryogenesis coupling mechanical behaviour and gene regulation. Nat Commun 8, 13929. 10.1038/ncomms13929

Dunn, S.-J., Näthke, I.S., Osborne, J.M., 2013. Computational Models Reveal a Passive Mechanism for Cell Migration in the Crypt. PLOS ONE 8, e80516. 10.1371/journal.pone.0080516

Economou, A.D., Ohazama, A., Porntaveetus, T., Sharpe, P.T., Kondo, S., Basson, M.A., Gritli-Linde, A., Cobourne, M.T., Green, J.B.A., 2012. Periodic stripe formation by a Turing mechanism operating at growth zones in the mammalian palate. Nat Genet 44, 348–351. 10.1038/ng.1090

Eldred, M.K., Charlton-Perkins, M., Muresan, L., Harris, W.A., 2017. Self-organising aggregates of zebrafish retinal cells for investigating mechanisms of neural lamination. Development dev.142760. 10.1242/dev.142760

Emmons-Bell, M., Durant, F., Hammelman, J., Bessonov, N., Volpert, V., Morokuma, J., Pinet, K., Adams, D., Pietak, A., Lobo, D., Levin, M., 2015. Gap Junctional Blockade Stochastically Induces Different Species-Specific Head Anatomies in Genetically Wild-Type Girardia dorotocephala Flatworms. International Journal of Molecular Sciences 16, 27865–27896. 10.3390/ijms161126065

Farahani, P.E., Nelson, C.M., 2022. Revealing epithelial morphogenetic mechanisms through live imaging. Current Opinion in Genetics & Development 72, 61–68. 10.1016/j.gde.2021.10.007

George, A., Akhavan, N., Peercy, B.E., Starz-Gaiano, M., 2025. Chemotaxis of Drosophila border cells is modulated by tissue geometry through dispersion of chemoattractants. iScience 28, 111959. 10.1016/j.isci.2025.111959

Germann, P., Marin-Riera, M., Sharpe, J., 2019. ya||a: GPU-Powered Spheroid Models for Mesenchyme and Epithelium. Cell Systems 8, 261–266.e3. 10.1016/j.cels.2019.02.007

Ghaffarizadeh, A., Heiland, R., Friedman, S.H., Mumenthaler, S.M., Macklin, P., 2018. PhysiCell: An open source physics-based cell simulator for 3-D multicellular systems. PLOS Computational Biology 14, e1005991. 10.1371/journal.pcbi.1005991

Gierer, A., Meinhardt, H., 1972. A theory of biological pattern formation. Kybernetik 12, 30–39. 10.1007/BF00289234

Glen, C.M., Kemp, M.L., Voit, E.O., 2019. Agent-based modeling of morphogenetic systems: Advantages and challenges. PLoS Comput Biol 15, e1006577. 10.1371/journal.pcbi.1006577

Green, J.B. a., Sharpe, J., 2015. Positional information and reaction-diffusion: two big ideas in developmental biology combine. Development 142, 1203–1211. 10.1242/dev.114991

Groves, I., Placzek, M., Fletcher, A.G., 2020. Of mitogens and morphogens: modelling Sonic Hedgehog mechanisms in vertebrate development. Phil. Trans. R. Soc. B 375, 20190660. 10.1098/rstb.2019.0660

Gurley, K.A., Rink, J.C., Alvarado, A.S., 2008a. β-Catenin Defines Head Versus Tail Identity During Planarian Regeneration and Homeostasis. Science 319, 323–327. 10.1126/science.1150029

Gurley, K.A., Rink, J.C., Sánchez Alvarado, A., Alvarado, A.S., 2008b. β-Catenin defines head versus tail identity during planarian regeneration and homeostasis. Science 319, 323–327. 10.1126/science.1150029

Hagolani, P.F., Zimm, R., Marin-Riera, M., Salazar-Ciudad, I., 2019. Cell signaling stabilizes morphogenesis against noise. Development 146, dev179309. 10.1242/dev.179309

Heisenberg, C.-P., Bellaïche, Y., 2013. Forces in Tissue Morphogenesis and Patterning. Cell 153, 948–962. 10.1016/j.cell.2013.05.008

Herath, S., Lobo, D., 2020. Cross-inhibition of Turing patterns explains the self-organized regulatory mechanism of planarian fission. Journal of Theoretical Biology 485, 110042. 10.1016/j.jtbi.2019.110042

Hill, E.M., Petersen, C.P., 2015. Wnt/Notum spatial feedback inhibition controls neoblast differentiation to regulate reversible growth of the planarian brain. Development 142, 4217–4229. 10.1242/dev.123612

Hopyan, S., Sharpe, J., Yang, Y., 2011. Budding behaviors: Growth of the limb as a model of morphogenesis. Dev. Dyn. 240, 1054–1062. 10.1002/dvdy.22601

Jung, H.-S., Francis-West, P.H., Widelitz, R.B., Jiang, T.-X., Ting-Berreth, S., Tickle, C., Wolpert, L., Chuong, C.-M., 1998. Local Inhibitory Action of BMPs and Their Relationships with Activators in Feather Formation: Implications for Periodic Patterning. Developmental Biology 196, 11–23. 10.1006/dbio.1998.8850

Kaul, H., Werschler, N., Jones, R.D., Siu, M.M., Tewary, M., Hagner, A., Ostblom, J., Aguilar-Hidalgo, D., Zandstra, P.W., 2023. Virtual cells in a virtual microenvironment recapitulate early development-like patterns in human pluripotent stem cell colonies. Stem Cell Reports 18, 377–393. 10.1016/j.stemcr.2022.10.004

Keller, R., 2012. Physical Biology Returns to Morphogenesis. Science 338, 201–203. 10.1126/science.1230718

Kicheva, A., Briscoe, J., 2023. Control of Tissue Development by Morphogens. Annual Review of Cell and Developmental Biology 39. 10.1146/annurev-cellbio-020823-011522

Ko, J.M., Lobo, D., 2019. Continuous Dynamic Modeling of Regulated Cell Adhesion: Sorting, Intercalation, and Involution. Biophysical Journal 117, 2166–2179. 10.1016/j.bpj.2019.10.032

Ko, J.M., Mousavi, R., Lobo, D., 2022. Computational Systems Biology of Morphogenesis, in: Cortassa, S., Aon, M.A. (Eds.), Computational Systems Biology in Medicine and Biotechnology: Methods and Protocols, Methods in Molecular Biology. Springer US, New York, NY, pp. 343–365. 10.1007/978-1-0716-1831-8_14

Ko, J.M., Reginato, W., Wolff, A., Lobo, D., 2024. Mechanistic regulation of planarian shape during growth and degrowth. Development 151, dev202353. 10.1242/dev.202353

Koch, A.J., Meinhardt, H., 1994. Biological pattern formation: from basic mechanisms to complex structures. Rev. Mod. Phys. 66, 1481–1507. 10.1103/RevModPhys.66.1481

Krause, A.L., Gaffney, E.A., Walker, B.J., 2023. Concentration-Dependent Domain Evolution in Reaction–Diffusion Systems. Bull Math Biol 85, 14. 10.1007/s11538-022-01115-2

Kudla, A.M., Miranda, X., Nijhout, H.F., 2022. The roles of growth regulation and appendage patterning genes in the morphogenesis of treehopper pronota. Proceedings of the Royal Society B: Biological Sciences 289, 20212682. 10.1098/rspb.2021.2682

Law, T.R., Hancox, J., Wright, S.A., Jarvis, S.A., 2019. An algorithm for computing short-range forces in molecular dynamics simulations with non-uniform particle densities. Journal of Parallel and Distributed Computing 130, 1–11. 10.1016/j.jpdc.2019.03.008

Lobo, D., 2022. Formalizing Phenotypes of Regeneration, in: Blanchoud, S., Galliot, B. (Eds.), Whole-Body Regeneration: Methods and Protocols, Methods in Molecular Biology. Springer US, New York, NY, pp. 663–679. 10.1007/978-1-0716-2172-1_36

Lobo, D., Beane, W.S., Levin, M., 2012. Modeling Planarian Regeneration: A Primer for Reverse-Engineering the Worm. PLoS Comput Biol 8, e1002481. 10.1371/journal.pcbi.1002481

Lobo, D., Levin, M., 2015. Inferring regulatory networks from experimental morphological phenotypes: a computational method reverse-engineers planarian regeneration. PLoS Computational Biology 11, e1004295. 10.1371/journal.pcbi.1004295

Lobo, D., Malone, T.J., Levin, M., 2013. Planform: an application and database of graph-encoded planarian regenerative experiments. Bioinformatics 29, 1098–1100. 10.1093/bioinformatics/btt088

Lu, P., Minowada, G., Martin, G.R., 2006. Increasing *Fgf4* expression in the mouse limb bud causes polysyndactyly and rescues the skeletal defects that result from loss of *Fgf8* function. Development 133, 33–42. 10.1242/dev.02172

Maini, P.K., Woolley, T.E., Baker, R.E., Gaffney, E.A., Lee, S.S., 2012. Turing’s model for biological pattern formation and the robustness problem. Interface Focus. 2, 487–496. 10.1098/rsfs.2011.0113

Marcon, L., Sharpe, J., 2012. Turing patterns in development: what about the horse part? Current Opinion in Genetics & Development 22, 578–584. 10.1016/j.gde.2012.11.013

Marin-Riera, M., Brun-Usan, M., Zimm, R., V??likangas, T., Salazar-Ciudad, I., 2015. Computational modeling of development by epithelia, mesenchyme and their interactions: A unified model. Bioinformatics 32, 219–225. 10.1093/bioinformatics/btv527

Maroudas-Sacks, Y., Suganthan, S., Garion, L., Ascoli-Abbina, Y., Westfried, A., Dori, N., Pasvinter, I., Popović, M., Keren, K., 2025. Mechanical strain focusing at topological defect sites in regenerating Hydra. Development 152, DEV204514. 10.1242/dev.204514

Mateus, R., Fuhrmann, J.F., Dye, N.A., 2021. Growth across scales: Dynamic signaling impacts tissue size and shape. Current Opinion in Cell Biology, Differentiation and development 73, 50–57. 10.1016/j.ceb.2021.05.002

Mathias, S., Coulier, A., Bouchnita, A., Hellander, A., 2020. Impact of Force Function Formulations on the Numerical Simulation of Centre-Based Models. Bull Math Biol 82, 132. 10.1007/s11538-020-00810-2

Mathias, S., Coulier, A., Hellander, A., 2022. CBMOS: a GPU-enabled Python framework for the numerical study of center-based models. BMC Bioinformatics 23, 55. 10.1186/s12859-022-04575-4

Mousavi, R., Konuru, S.H., Lobo, D., 2021. Inference of dynamic spatial GRN models with multi-GPU evolutionary computation. Briefings in Bioinformatics 22, 1–11. 10.1093/bib/bbab104

Mousavi, R., Lobo, D., 2024. Automatic design of gene regulatory mechanisms for spatial pattern formation. npj Syst Biol Appl 10, 1–13. 10.1038/s41540-024-00361-5

Moustakas-Verho, J.E., Zimm, R., Cebra-Thomas, J., Lempiäinen, N.K., Kallonen, A., Mitchell, K.L., Hämäläinen, K., Salazar-Ciudad, I., Jernvall, J., Gilbert, S.F., 2014. The origin and loss of periodic patterning in the turtle shell. Development 141, 3033–3039. 10.1242/dev.109041

Muñoz-Nava, L.M., Alvarez, H.A., Flores-Flores, M., Chara, O., Nahmad, M., 2020. A dynamic cell recruitment process drives growth of the Drosophila wing by overscaling the vestigial expression pattern. Developmental Biology 462, 141–151. 10.1016/j.ydbio.2020.03.009

Okuda, S., Miura, T., Inoue, Y., Adachi, T., Eiraku, M., 2018a. Combining Turing and 3D vertex models reproduces autonomous multicellular morphogenesis with undulation, tubulation, and branching. Sci Rep 8, 2386. 10.1038/s41598-018-20678-6

Okuda, S., Miura, T., Inoue, Y., Adachi, T., Eiraku, M., 2018b. Combining Turing and 3D vertex models reproduces autonomous multicellular morphogenesis with undulation, tubulation, and branching. Sci Rep 8, 2386. 10.1038/s41598-018-20678-6

Osborne, J.M., Fletcher, A.G., Pitt-Francis, J.M., Maini, P.K., Gavaghan, D.J., 2017. Comparing individual-based approaches to modelling the self-organization of multicellular tissues. PLoS Computational Biology 13, e1005387. 10.1371/journal.pcbi.1005387

Pleyer, J., Fleck, C., 2023. Agent-based models in cellular systems. Front. Phys. 10. 10.3389/fphy.2022.968409

Purcell, E.M., 1977. Life at low Reynolds number. American Journal of Physics 45, 3–11. 10.1119/1.10903

Ramezani, A., Britton, S., Zandi, R., Alber, M., Nematbakhsh, A., Chen, W., 2023. A multiscale chemical-mechanical model predicts impact of morphogen spreading on tissue growth. npj Syst Biol Appl 9, 1–12. 10.1038/s41540-023-00278-5

Raspopovic, J., Marcon, L., Russo, L., Sharpe, J., 2014. Digit patterning is controlled by a Bmp-Sox9-Wnt Turing network modulated by morphogen gradients. Science 345, 566–570. 10.1126/science.1252960

Reddien, P.W., 2021. Positional Information and Stem Cells Combine to Result in Planarian Regeneration. Cold Spring Harbor Perspectives in Biology a040717. 10.1101/cshperspect.a040717

Reddien, P.W., 2018. The Cellular and Molecular Basis for Planarian Regeneration. Cell 175, 327–345. 10.1016/j.cell.2018.09.021

Roy, J., Cheung, E., Bhatti, J., Muneem, A., Lobo, D., 2020. Curation and annotation of planarian gene expression patterns with segmented reference morphologies. Bioinformatics 36, 2881–2887. 10.1093/bioinformatics/btaa023

Runser, S., Vetter, R., Iber, D., 2024. SimuCell3D: three-dimensional simulation of tissue mechanics with cell polarization. Nat Comput Sci 4, 299–309. 10.1038/s43588-024-00620-9

Salbreux, G., Barthel, L.K., Raymond, P.A., Lubensky, D.K., 2012. Coupling Mechanical Deformations and Planar Cell Polarity to Create Regular Patterns in the Zebrafish Retina. PLoS Comput Biol 8, e1002618. 10.1371/journal.pcbi.1002618

Salm, M., Pismen, L.M., 2012. Chemical and mechanical signaling in epithelial spreading. Phys. Biol. 9, 026009. 10.1088/1478-3975/9/2/026009

Schnakenberg, J., 1979. Simple chemical reaction systems with limit cycle behaviour. Journal of Theoretical Biology 81, 389–400. 10.1016/0022-5193(79)90042-0

Scimone, M.L., Cote, L.E., Rogers, T., Reddien, P.W., 2016. Two FGFRL-Wnt circuits organize the planarian anteroposterior axis. eLife 5, e12845. 10.7554/eLife.12845

Sharpe, J., 2017. Computer modeling in developmental biology: growing today, essential tomorrow. Development 144, 4214–4225. 10.1242/dev.151274

Shu, W., Kaplan, C.N., 2023. A multiscale whole-cell theory for mechanosensitive migration on viscoelastic substrates. Biophysical Journal 122, 114–129. 10.1016/j.bpj.2022.11.022

Steinberg, M.S., 1958. On the Chemical Bonds between Animal Cells. A Mechanism for Type-Specific Association. The American Naturalist 92, 65–81. 10.1086/282013

Stückemann, T., Cleland, J.P., Werner, S., Thi-Kim Vu, H., Bayersdorf, R., Liu, S.-Y., Friedrich, B., Jülicher, F., Rink, J.C., 2017. Antagonistic Self-Organizing Patterning Systems Control Maintenance and Regeneration of the Anteroposterior Axis in Planarians. Developmental Cell 40, 248–263.e4. 10.1016/j.devcel.2016.12.024

Sureda-Gómez, M., Martín-Durán, J.M., Adell, T., 2016. Localization of planarian β-CATENIN-1 reveals multiple roles during anterior-posterior regeneration and organogenesis. Development 143, 4149–4160. 10.1242/dev.135152

Thompson, K.W., Joshi, P., Dymond, J.S., Gorrepati, L., Smith, H.E., Krause, M.W., Eisenmann, D.M., 2016. The Paired-box protein PAX-3 regulates the choice between lateral and ventral epidermal cell fates in *C. elegans*. Developmental Biology 412, 191–207. 10.1016/j.ydbio.2016.03.002

Townes, P.L., Holtfreter, J., 1955. Directed movements and selective adhesion of embryonic amphibian cells. J. Exp. Zool. 128, 53–120. 10.1002/jez.1401280105

Truskey, G.A., Yuan, F., Katz, D.F., 2009. Transport phenomena in biological systems, 2nd ed. ed, Pearson Prentice Hall bioengineering. Pearson Prentice Hall, Upper Saddle River, N.J.

Van Leeuwen, I.M.M., Mirams, G.R., Walter, A., Fletcher, A., Murray, P., Osborne, J., Varma, S., Young, S.J., Cooper, J., Doyle, B., Pitt-Francis, J., Momtahan, L., Pathmanathan, P., Whiteley, J.P., Chapman, S.J., Gavaghan, D.J., Jensen, O.E., King, J.R., Maini, P.K., Waters, S.L., Byrne, H.M., 2009. An integrative computational model for intestinal tissue renewal. Cell Proliferation 42, 617–636. 10.1111/j.1365-2184.2009.00627.x

Van Liedekerke, P., Palm, M.M., Jagiella, N., Drasdo, D., 2015. Simulating tissue mechanics with agent-based models: concepts, perspectives and some novel results. Comp. Part. Mech. 2, 401–444. 10.1007/s40571-015-0082-3

Vasilopoulos, G., Painter, K.J., 2016. Pattern formation in discrete cell tissues under long range filopodia-based direct cell to cell contact. Mathematical Biosciences 273, 1–15. 10.1016/j.mbs.2015.12.008

Vazquez, K., Saraswathibhatla, A., Notbohm, J., 2022. Effect of substrate stiffness on friction in collective cell migration. Sci Rep 12, 2474. 10.1038/s41598-022-06504-0

Werner, S., Stückemann, T., Beirán Amigo, M., Rink, J.C., Jülicher, F., Friedrich, B.M., Amigo, M.B., Rink, J.C., Jülicher, F., Friedrich, B.M., 2015. Scaling and regeneration of self-organized patterns. Physical review letters 114, 138101. 10.1103/PhysRevLett.114.138101

West, J., Robertson-Tessi, M., Anderson, A.R.A., 2023. Agent-based methods facilitate integrative science in cancer. Trends in Cell Biology 33, 300–311. 10.1016/j.tcb.2022.10.006

Witchley, J.N., Mayer, M., Wagner, D.E., Owen, J.H., Reddien, P.W., 2013. Muscle Cells Provide Instructions for Planarian Regeneration. Cell Reports 4, 633–641. 10.1016/j.celrep.2013.07.022

Wolff, A., Wagner, C., Wolf, J., Lobo, D., 2022. In situ probe and inhibitory RNA synthesis using streamlined gene cloning with Gibson assembly. STAR Protocols 3, 101458. 10.1016/j.xpro.2022.101458

Xue, S.-L., Yang, Q., Liberali, P., Hannezo, E., 2025. Mechanochemical bistability of intestinal organoids enables robust morphogenesis. Nat. Phys. 21, 608–617. 10.1038/s41567-025-02792-1

Yamaguchi, M., Yoshimoto, E., Kondo, S., 2007. Pattern regulation in the stripe of zebrafish suggests an underlying dynamic and autonomous mechanism. Proc. Natl. Acad. Sci. U.S.A. 104, 4790–4793. 10.1073/pnas.0607790104

